# Direct interhemispheric cortical communication via thalamic commissures: a new white-matter pathway in the primate brain

**DOI:** 10.1101/2023.06.15.545128

**Authors:** Diego Szczupak, David J. Schaeffer, Xiaoguang Tian, Sang-Ho Choi, Fang-Cheng, Pamela Meneses Iack, Vinicius P. Campos, J. Patrick Mayo, Janina Patsch, Christian Mitter, Amit Haboosheh, Marcelo A.C. Vieira, Gregor Kasprian, Fernanda Tovar-Moll, Roberto Lent, Afonso C. Silva

## Abstract

Cortical neurons of eutherian mammals project to the contralateral hemisphere, crossing the midline primarily via the corpus callosum and the anterior, posterior, and hippocampal commissures. We recently reported an additional commissural pathway in rodents, termed the thalamic commissures (TCs), as another interhemispheric axonal fiber pathway that connects cortex to the contralateral thalamus. Here, we demonstrate that TCs also exist in primates and characterize the connectivity of these pathways with high-resolution diffusion-weighted magnetic resonance imaging, viral axonal tracing, and functional MRI. We present evidence of TCs in both New World (*Callithrix jacchus* and *Cebus apella*) and Old World primates (*Macaca mulatta*). Further, like rodents, we show that the TCs in primates develop during the embryonic period, forming anatomical and functionally active connections of the cortex with the contralateral thalamus. We also searched for TCs in the human brain, showing their presence in humans with brain malformations, although we could not identify TCs in healthy subjects. These results pose the TCs as an important fiber pathway in the primate brain, allowing for more robust interhemispheric connectivity and synchrony and serving as an alternative commissural route in developmental brain malformations.

**Significance statement:** Brain connectivity is a central topic in neuroscience. Understanding how brain areas can communicate allows for the comprehension of brain structure and function. We have described in rodents a new commissure pathway that connects the cortex to the contralateral thalamus. Here, we investigate whether this pathway exists in non-human primates and humans. The presence of these commissures poses the TCs as an important fiber pathway in the primate brain, allowing for more robust interhemispheric connectivity and synchrony and serving as an alternative commissural route in developmental brain malformations.

## Introduction

The eutherian mammalian brain has evolved to cluster neurons in specific brain regions that perform distinct functions, interconnected so that the whole is greater than the sum of its individual parts (1). These connections can be categorized as intrahemispheric, occurring within a cerebral hemisphere, or interhemispheric, crossing the midline through specific pathways. The corpus callosum (CC) is the main interhemispheric pathway in the Eutherian mammalian brain, followed by the anterior, posterior, and hippocampal commissures (2, 3).

Recently, we discovered a novel interhemispheric axonal fiber pathway in rodents connecting the cortex with the contralateral thalamus, which we named the thalamic commissures (TCs), formed by four distinct patches traversing the thalamus midline (4). These findings support previous reports in the literature that documented sporadic bilateral projections from the cortex to the contralateral thalamus, particularly from the hippocampus (5), prefrontal cortex (4, 6), and primary motor cortex (4, 7). This suggests that TCs may represent a normotypical pathway for interhemispheric connectivity and could be present in other high-order species.

In our previous work, we also found that corpus callosum dysgenesis (CCD), a developmental callosal malformation, altered the TC’s anatomical presentation and tissue properties (e.g., fractional anisotropy – FA) (4). Developmental malformations can generally result in anatomical alterations that disrupt the brain’s wiring. For example, CCD can lead to atypical anatomical white matter rewiring, such as the Probst bundle (8), sigmoid bundle (9–11), and intercortical bundle (12). Furthermore, these alterations lead to global network changes in human subjects (13–16) and have also been observed in non-human primates (17) and mice (18–20).

In this study, we sought to probe the existence of TCs in other mammalian species. We employed high-resolution diffusion imaging, viral axonal tracing, and fMRI techniques to investigate the TCs in three non-human primate species and comprehensively describe their macroscopic anatomy, fine axonal connectivity, development, and function. Furthermore, to extend the scope of our findings, we applied diffusion-weighted imaging to examine the presence of TCs in humans in both healthy human subjects and patients with brain malformations.

## Results

### Thalamic commissures in the non-human primate brain

We imaged the brains of three different non-human primates (NHP) species *ex vivo* with high-resolution DWI to look for TCs (Figure 1). We identified white matter tracts traversing the thalamus at the midline in all three monkey brains, suggesting TCs are a common feature of the primate brain. These TCs showed different presentations. In the marmoset, the TCs were primarily oriented diagonally along the anteroposterior axis and segmented into different elongated patches (Figure 1A). In the capuchin brain, the TCs were dispersed in a leminiscal fashion and split into two curved but parallel pathways along the inferior portions of the thalamus (Figure 1B). In the Rhesus macaque, we identified several patches of white matter fibers traversing the thalamus Figure 1C). We also identified TCs in marmosets in DWI scans acquired in vivo (Supplementary Figure 1), and in ultra-high-resolution ex-vivo T2* images (Supplementary Figure 2). These MRI modalities establish the presence of white matter fibers traversing the midline at the thalamus, allowing further investigation of the TCs.

**Figure 1.**
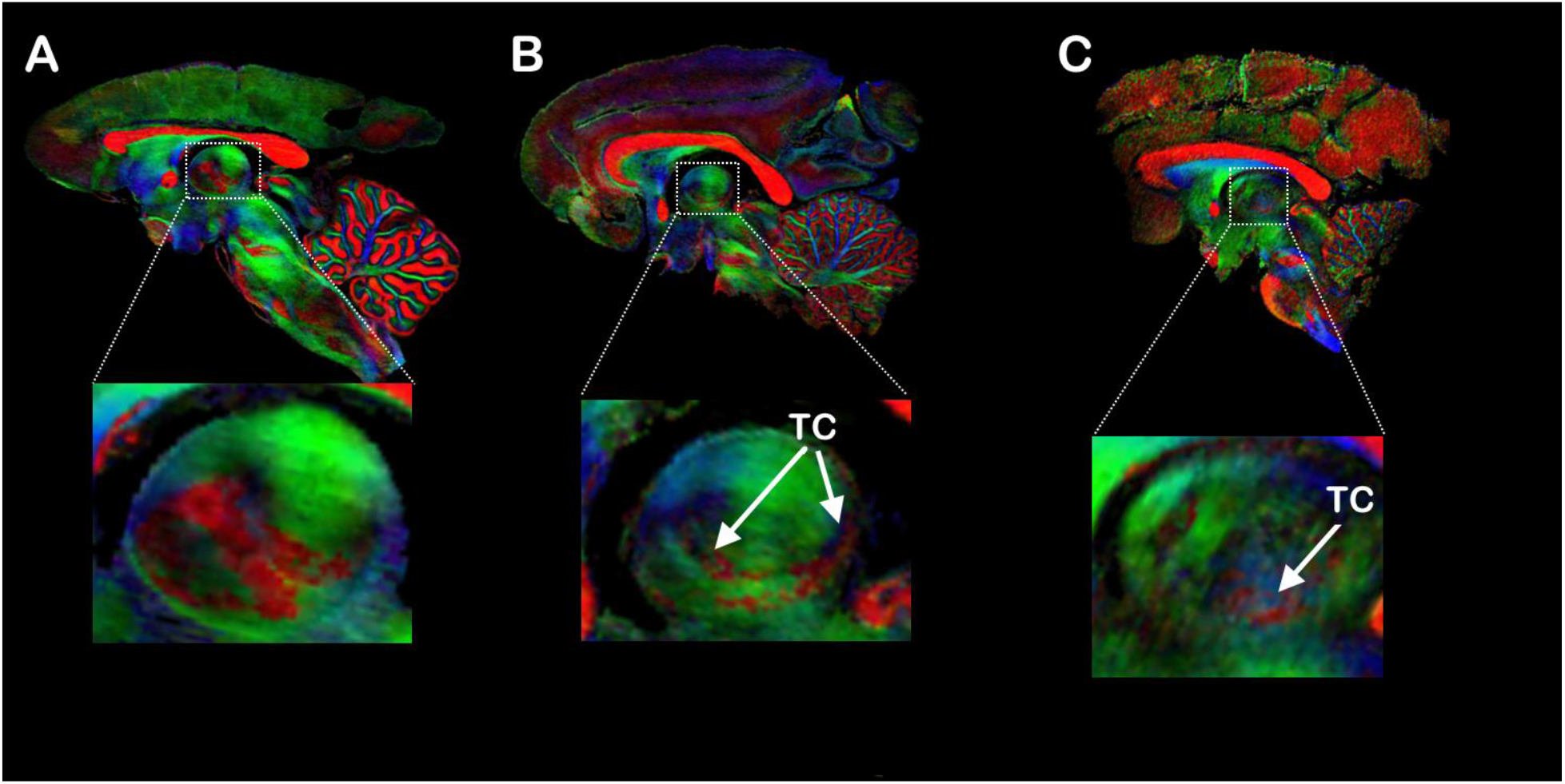
Non-human primate comparative thalamic commissures. Diffusion-encoded colormap (DEC) images of fractional anisotropy (FA) of a midsagittal brain slice of three different NHP species: **(A)** Marmoset, **(B)** Capuchin, and **(C)** Macaque. Insets show the presence of white matter fibers traversing the midline at the thalamus, suggesting the presence of Thalamic Commissures (TC, arrows). In the DEC images, red represents the mediolateral (ML), green represents the anteroposterior (AP), and blue represents the dorsoventral (DV) directions.

### TC axonal tracing in the marmoset brain

To validate the DWI results shown in Figure 1, we injected the brains of two marmosets with AAV9 anterograde tracer conjugated with GFP or Td-Tomato in two distinct brain regions. In one animal, the viral tracer was injected into the dorsolateral prefrontal cortex (A6DR - Figure 2A), an area analogous to the mouse lateral orbital cortex, that was recently described to connect with the contralateral hemisphere via the CC and TC (4, 6). Our data clearly show cohesive white matter fiber bundles crossing the midline via the CC (Figure 2B), and diffuse fibers traversing the midline via the TC (Figure 2 C-D). The second animal was injected in the hippocampus, a region that connects to the contralateral thalamus in macaques (5) via the reuniens nuclei. We injected the hippocampus in two different locations, one dorsal (Figure 2E, red) and the other ventral (Figure 2E, green). Our data show that the dorsal portion of the hippocampus communicates with the contralateral hemisphere through the superior colliculus (SC - Figure 2F) and the reuniens nuclei of the thalamus (Figure 2G-H). In contrast, fibers in the ventral portion of the hippocampus project to the contralateral hemisphere via the hippocampal commissure and the fornix (Fx, Figure 2G).

**Figure 2.**
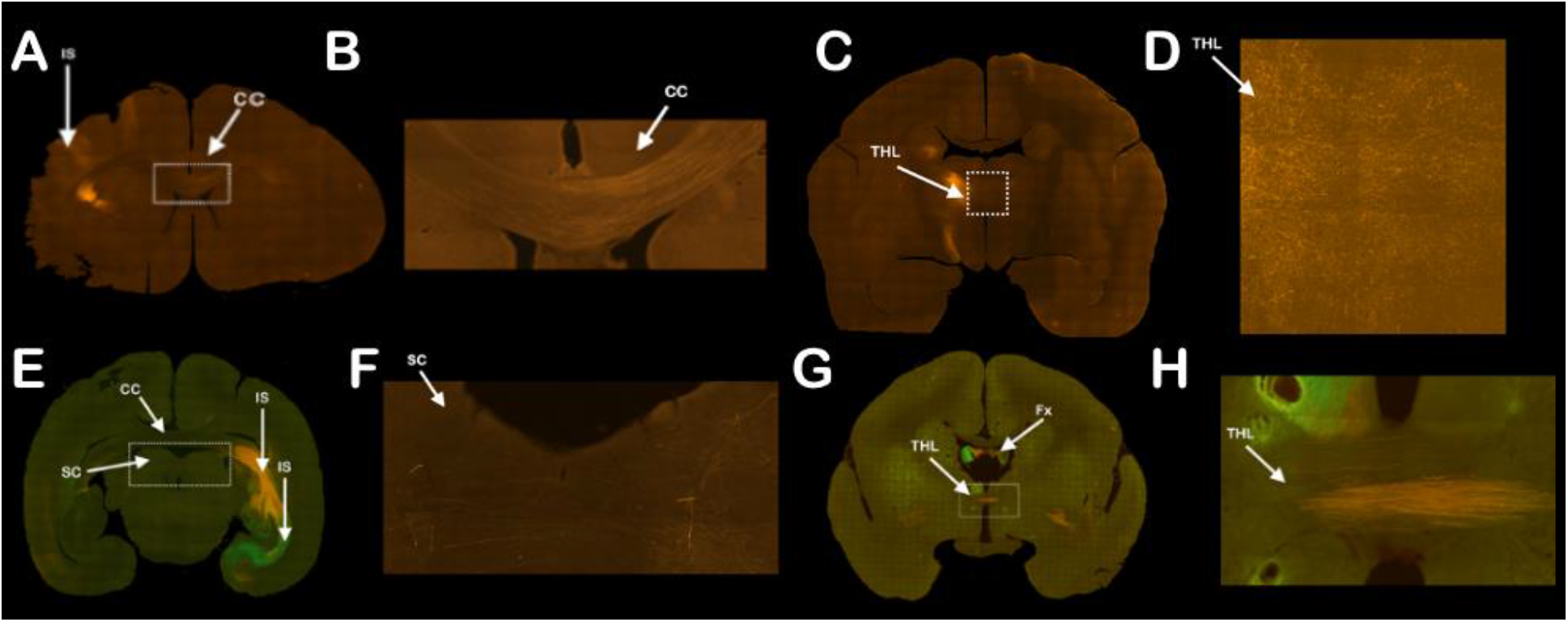
Viral axonal tracing of Thalamic commissures in the marmoset brain. Injection of AAV9 in two different cortical sites, the Dorsolateral prefrontal cortex **(A)** and Hippocampus **(E)**, to probe the interhemispheric connectivity of the TCs. The dorsolateral prefrontal cortex injection (red) revealed robust interhemispheric connectivity through the corpus callosum (CC) and diffuse connectivity through the thalamus **(A-D)**. The hippocampus was injected in two different locations, the dorsal hippocampus (red) and the ventral hippocampus (green) **(E)**. The dorsal injection site revealed interhemispheric connectivity through the superior colliculus (SC) **(F)**, the hippocampal commissure/fornix (Fx), and the thalamus (THL) **(G-H)**. In contrast, the ventral portion of the hippocampus only crosses the midline via the hippocampal commissure/fornix.

### Thalamic commissures normal development

We employed high-resolution *ex vivo* DWI to investigate the TCs development in marmosets and mice by scanning the brain of neonate (p0, Figures 3A, 3C) and adult (Figure 3B, 3D) animals. We could clearly resolve TCs at both ages, suggesting TCs are present at early developmental stages. To prevent confounding factors related to the MRI scanner, diffusion scheme, sequence differences, or inter-specific differences, we compared the ratio of FA of the TCs with other well-studied commissures (CC and AC). In both species, we found no statistical differences in the FA ratio between neonates and adults (Figure 3E-F). However, marmosets presented higher TC/CC ratio than mice (p=0.0187 – Figure 3E), likely due to CC differences in FA between marmosets and mice.

**Figure 3.**
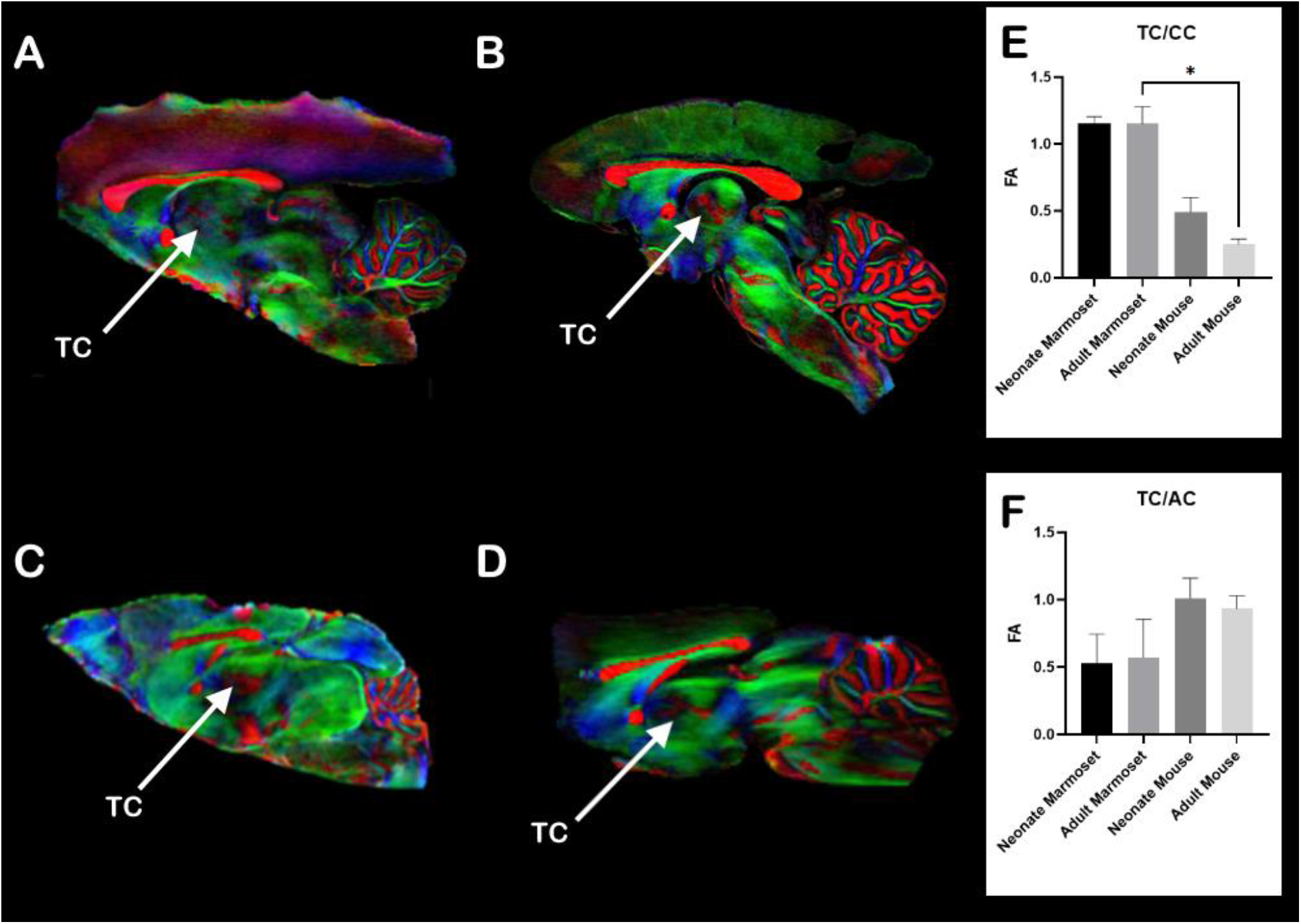
Thalamic commissures development. Direction**-**encoded diffusion-weighted images show the brain of a neonate **(A)** and an adult **(B)** marmoset and a neonate **(C)** and adult mouse **(D)** FA maps with the thalamic commissures indicated by the arrows. **(E)** A plot of the FA values of the thalamic commissures normalized by the genu of the corpus callosum (CC) and **(F)** the anterior commissure (AC). In the Direction-Encoded Colormaps, red represents mediolateral (ML) diffusion, green represents anteroposterior (AP) diffusion, and blue represents dorsoventral (DV) diffusion. *p-value = 0.0187.

### TC functional connectivity

To further evaluate the connectivity of brain areas with the contralateral thalamus, we investigated the functional connectivity populational averaged map of 31 adult marmosets (23). We analyzed the connectivity of a seed voxel within the dorsolateral prefrontal cortex (Figure 4A) and the hippocampus (Figure 4C) and verified that both of these regions were positively correlated to the contralateral thalamus (Figure 4B, 4D), indicating functional connectivity. Conversely, we placed a seed in the thalamus (Figure 4E) and observed both positive and negative correlation with cortical regions in the contralateral hemisphere (Figure 4F). The thalamic connectivity spreads throughout the contralateral hemisphere, posing the TCs as an essential connectivity hub across hemispheres.

**Figure 4.**
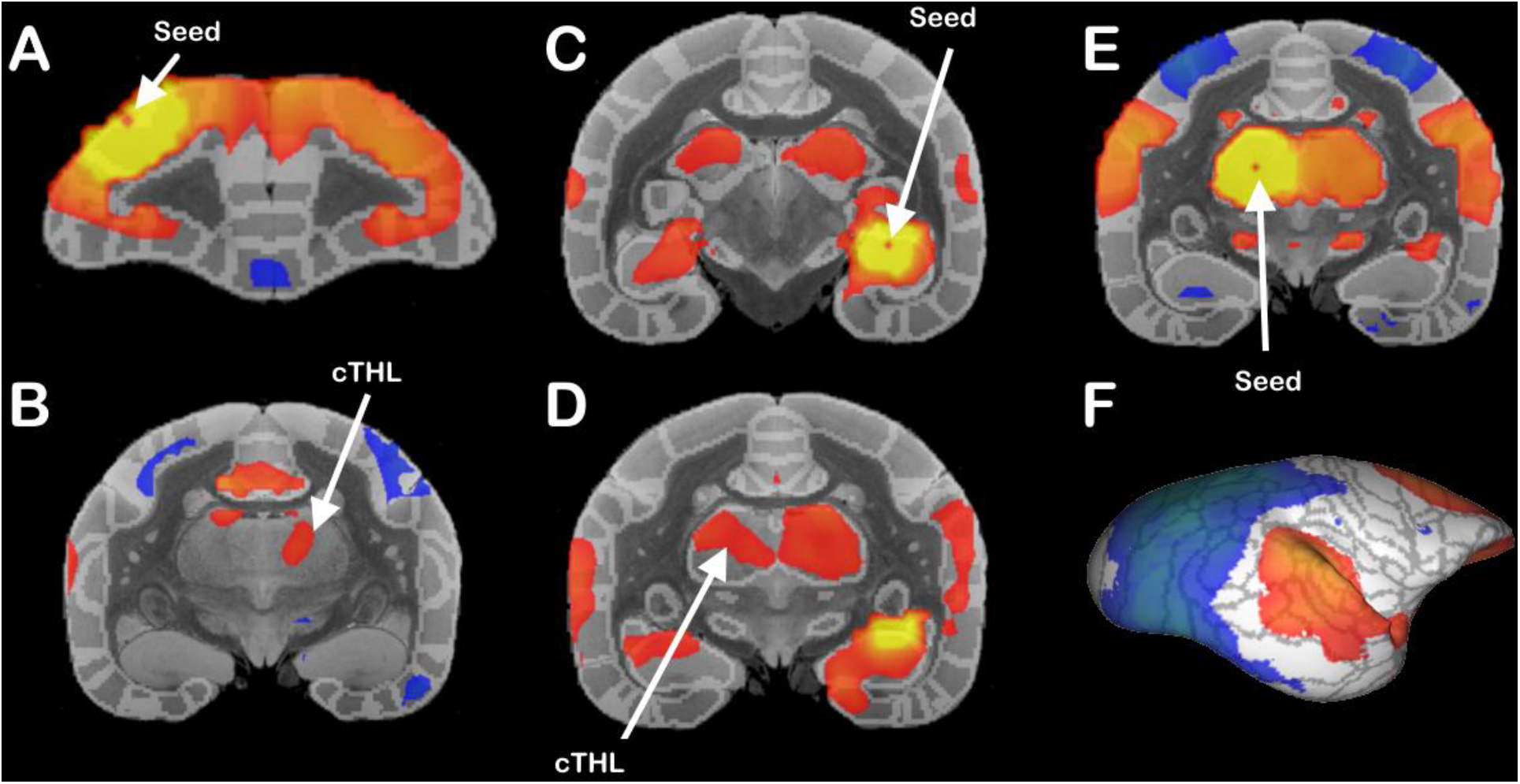
Marmoset functional connectivity. Functional resting-state connectivity averaged populational correlation map of 31 marmosets reveals cortical functional connectivity with the contralateral thalamus. We used single-voxel seeds in the same ROIS that we had previously used for histological tracing: the dorsolateral prefrontal cortex **(A)** and dorsal hippocampus **(C)** and show that both areas are functionally connected to the contralateral thalamus (cTHL) (**B** and **D** respectively). Thalamic seed **(E)** and contralateral cortical functional connectivity across the whole hemisphere **(F)**.

### Human thalamic commissures

In addition to identifying the TCs in the NHP brain, we investigated their presence in the human brain. In a super-resolution DWI scan of a healthy subject, we could not identify any evidence of TCs in sagittal, coronal, or axial views (Figure 5A-F). However, a callosal dysgenesis (CCD) patient clearly showed a TC (Figure 5G-L). In a cohort of 11 CCD patients, we identified TCs in 4 subjects (Supplementary Figure 3). We also identified TCs in two infants from an Austrian patient cohort with non-CCD brain malformations (Figure 6), indicating that the TCs may be associated with many different brain malformations. Our data show that, while not normally discernible in the healthy human brain, TCs can form in subjects afflicted with developmental disorders, posing the TCs as an alternative pathway for interhemispheric communication in primates.

**Figure 5.**
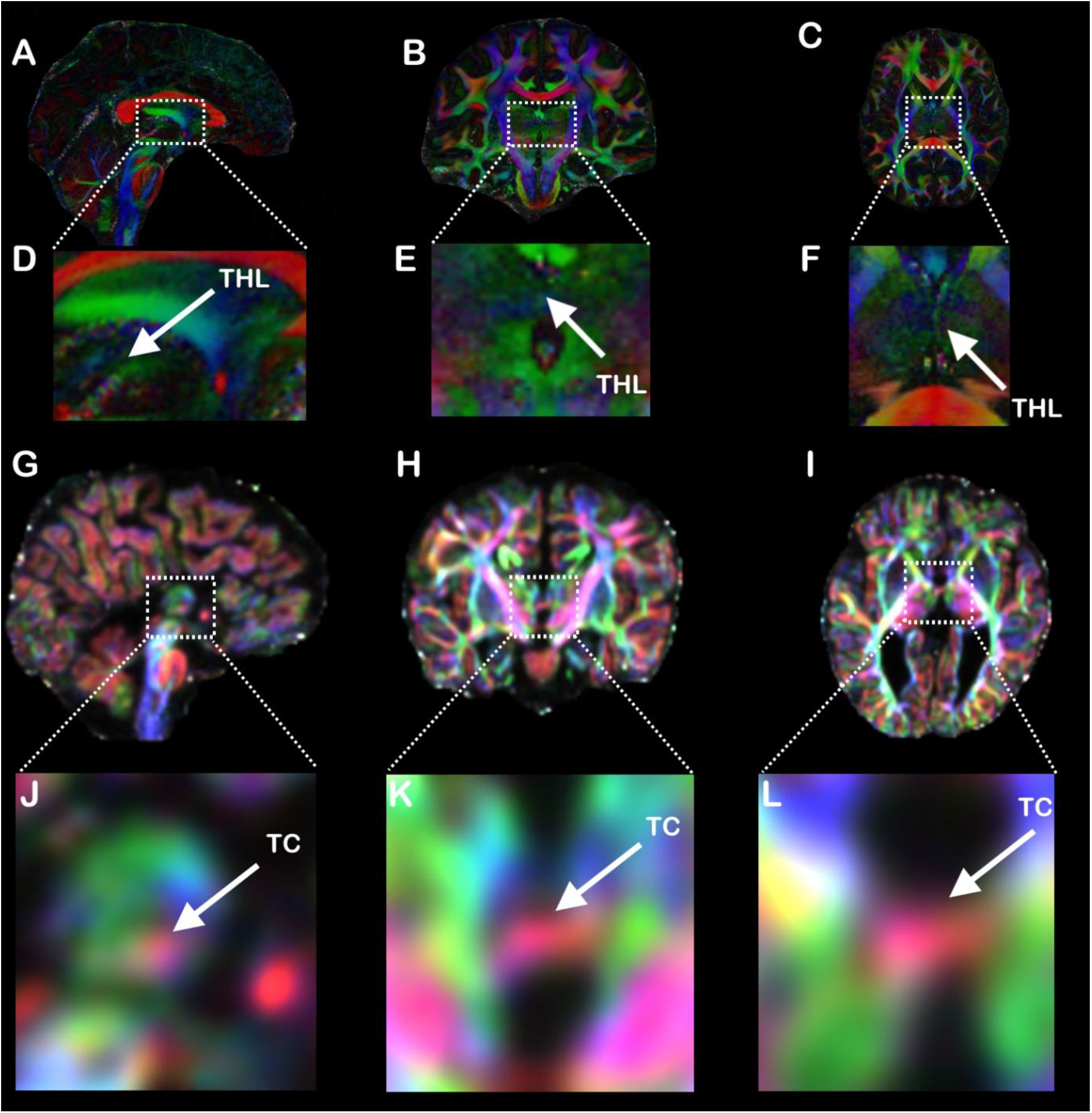
Human Thalamic commissures. Direction-Encoded Colormaps of diffusion-weighted FA showing the thalamic region of a healthy human subject in sagittal **(A)**, coronal **(B)**, and axial **(C)** views. No visible TCs were detected on the zoomed insets **(D-F)**. Equivalent brain views of a patient with callosal dysgenesis **(G-I)** showing a clear thalamic commissure on the insets **(J-l)**. In the Direction-Encoded Colormaps, red represents mediolateral (ML) diffusion, green represents anteroposterior (AP) diffusion, and blue represents dorsoventral (DV) diffusion.

**Figure 6.**
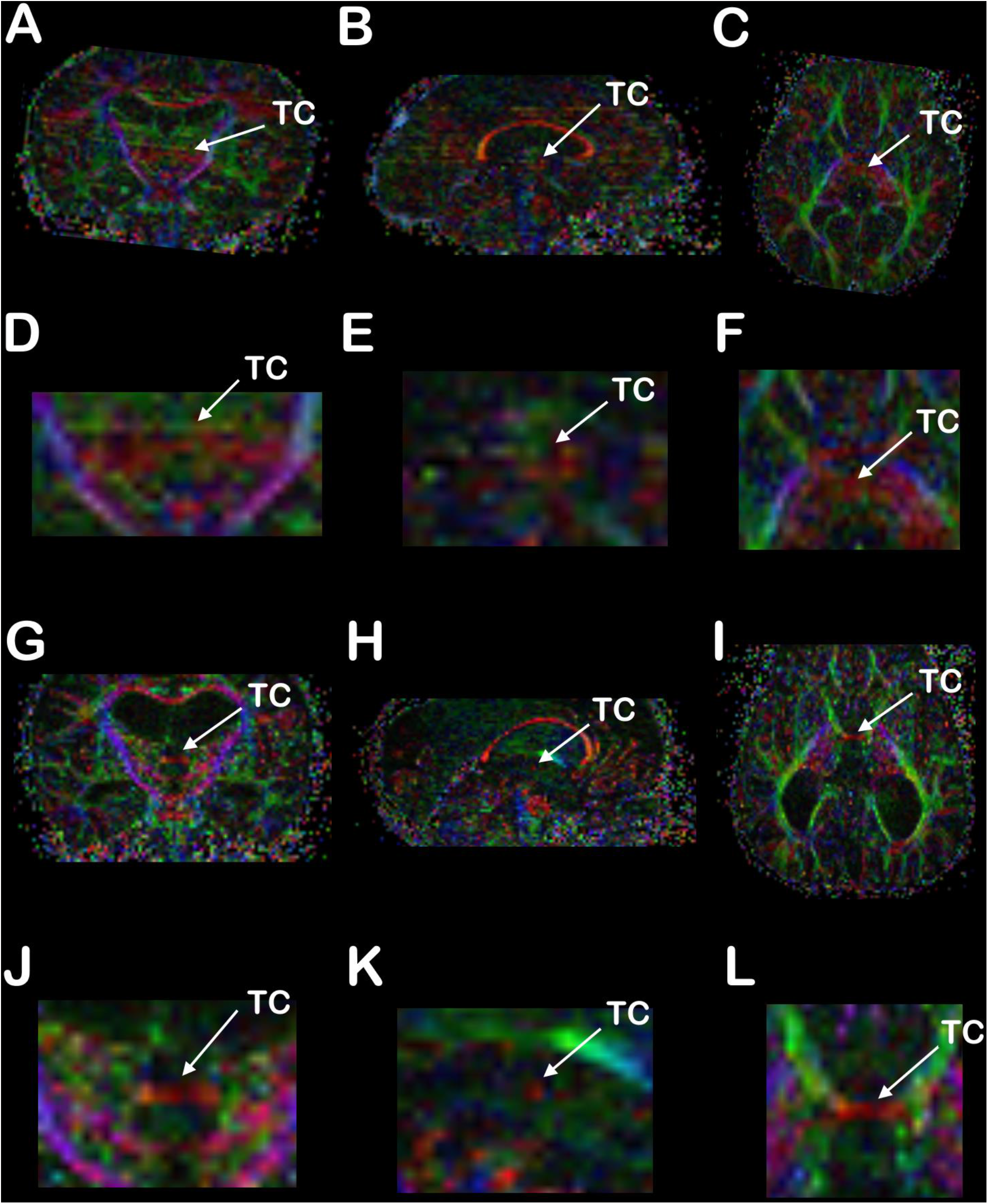
Human Infant Thalamic commissures. Direction-Encoded Colormaps in coronal **(A, G)**, sagittal **(B, H)**, and axial **(C, I)** views of diffusion-weighted FA showing the thalamic region of two infants (**A-F** and **G-L**) with brain malformations. Visible TCs were detected on the zoomed insets **(D-F and J-L)**. In the Direction-Encoded Colormaps, red represents mediolateral (ML) diffusion, green represents anteroposterior (AP) diffusion, and blue represents dorsoventral (DV) diffusion.

## Discussion

In this work, we leverage high-resolution diffusion-weighted imaging, viral axonal fiber tracking, and functional MRI to accurately describe and characterize the TCs in the primate brain at birth and adulthood. We verified the existence of TCs in three non-human primate species and humans with developmental callosal malformations. We found that the TCs are present in the neonate brain, suggesting that their morphogenesis follows the primary commissural system. Furthermore, we described how the TCs contribute to a clear functional connectivity of the thalamus with the contralateral cortex. Altogether, our findings pose the TCs as a common interhemispheric pathway in the callosal mammalian brain.

### TCs in the NHP brain

It is necessary to distinguish the TCs from the inter-thalamic adherence, an anatomical structure of grey and white matter defined by the union between the hemispheric sides of the thalamus at the midline. Here, we refer to TCs as organized white-matter fiber bundles connecting the cortex to the contralateral thalamus. In previous work, we identified the TCs in C57BL6 and Balb/c mice (4). The TCs in the C57BL6 mouse brain presented a stereotypical anatomical formation with four distinct patches distributed along the anteroposterior axis of the thalamus. Here, we confirmed that TCs also exist in the brains of three non-human primate species. However, unlike rodents, the anatomical organization of the TCs along the thalamic midline varied substantially across the three species. Marmosets presented a clear and well-organized TC composed of different elongated patches distributed diagonally along the thalamus, while the capuchin monkey’s TCs spread along two ribbon-like patterns, and the macaque showed a broader dispersal of TCs.

Although the TC is a central commissure connecting the brain hemispheres, it is distinct from the large AC of metatherian mammals, such as marsupials (2), which connects both cortical hemispheres in absence of the CC. It remains to be shown whether marsupials also present the TCs.

### Axonal connectivity

We used AAV-9 anterograde neuronal tracers to verify the results shown with diffusion-weighted imaging of the marmoset brain. Our marmoset hippocampal AAV-9 histological findings agree with previous macaque (5) and mice (4) findings that the hippocampus directly connects the contralateral thalamus. In addition, we used a dual injection of AAV-9 expressing fluorescent markers of distinct colors to show that only the dorsal hippocampus communicates via the TC, indicating the presence of two separate hippocampal circuits with distinct interhemispheric communications. This set of results adds to the pool of literature that the dorsal and ventral hippocampus are different functional regions (35).

Unlike the hippocampus’s compact connectivity, the AAV-9 injection in the dorsolateral prefrontal cortex showed dispersed connectivity to the contralateral thalamus via the TCs. This diffuse connectivity pattern helps us understand why these connections are difficult to map, as they are sparse in histology and have low FA, giving a faint signal in diffusion-weighted imaging. Additionally, they connect a large portion of the contralateral thalamus, thus influencing many contralateral cortical regions. The dorsolateral prefrontal cortex is a central hub of the default-mode network (DMN) (36); therefore, it is reasonable to link these commissures with the DMN and its interhemispheric synchrony.

### Development

It is well documented that white-matter fiber bundles change along development through life, into aging. Although most fibers are developed during the gestational period, myelination (37), pruning (38), and fine dendritic morphology(39) are plastic and vary longitudinally. Our work investigated brain changes with MRI across species and ages. Therefore, we compared the TCs FA normalized by the AC and CC FA to avoid confounding effects from the difference in pulse sequences or MRI scanners and hardware (40). We have found no differences between neonates and adults in both marmosets and mice, indicating that the TCs have a similar development as the CC and AC and are likely regulated by similar processes.

However, we have found a statistically significant difference between the marmoset adult TC/CC ratio relative to adult mice, but no difference in TC/AC ratio. Therefore, this is likely due to differences in the CC FA between species. This increase in CC FA agrees with a larger cortical volume in primates relative to rodents(41).

### Functional connectivity

After investigating the structural connectivity of the TCs in the NHP brain, we analyzed the functional connectivity of the marmoset brain to understand if these axonal fibers that cross the midline via the TC can functionally connect brain areas. As functional connectivity is a mix of monosynaptic and polysynaptic connections(42), these circuits might not be mediated only by the TCs.

Nevertheless, monosynaptic connections of the TCs contribute to net functional connectivity.

As the thalamus is functionally connected to the contralateral hemisphere, it is reasonable to assume that the thalamus is not only a mediator of ipsilateral cortical connectivity (43) but contralateral connectivity. Here, we hypothesize that the TCs (Figure 7, blue and purple) allow the thalamus to receive contralateral cortical information to facilitate or inhibit cortical regions that will receive contralateral cortical information via other commissures (Figure 7, magenta). Therefore, the thalamus is a control hub that mediates cortico-cortical interhemispheric connectivity and synchrony (Figure 7).

**Figure 7.**
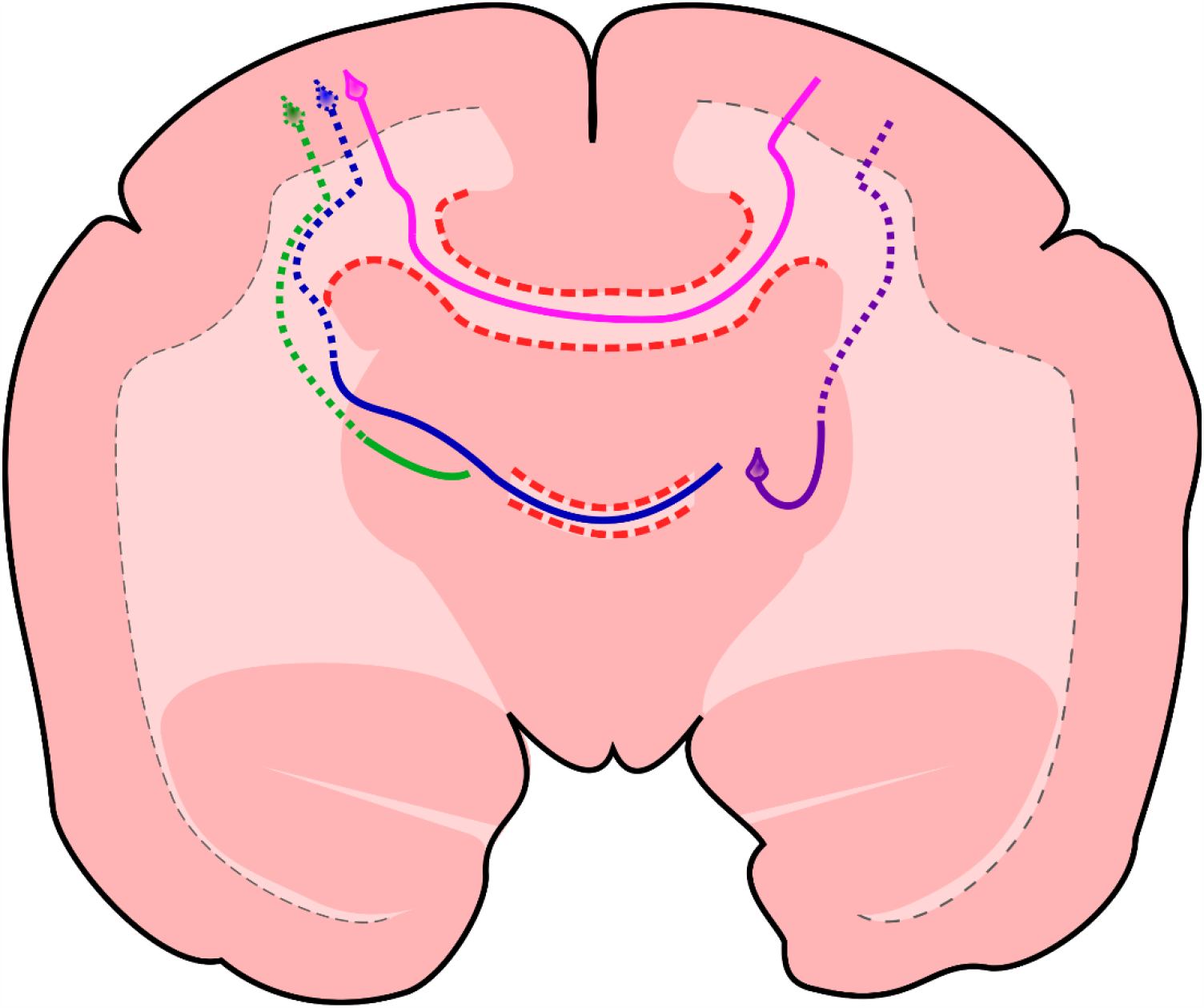
Schematic figure portraying the TCs in a marmoset brain. Coronal view of a marmoset brain showing the CC in magenta, intrahemispheric ipsilateral cortico-thalamic communication in green, intrahemispheric contralateral cortico-thalamic communication in purple, and the TCs in blue. This circuit allows the thalamus to communicate with its own hemisphere and play a role in interhemispheric cortical connectivity.

### Human TC

We used an ultra-high-resolution diffusion image to investigate the presence of the TCs in the healthy human brain. We could not find any white matter fibers that cross the midline via the thalamus similar to the TCs. This was expected as most human thalami are anatomically separated.

However, we identified the TCs in half of the CCD (with a higher prevalence of fused thalami) and on the Austrian brain malformation infant cohort. We hypothesize that the molecular and cellular substrate to the TCs development is present in the healthy human brain, but since the thalami are separated, the fibers cannot project to their natural target. When, for any reason, the thalami are unified, the TCs can form naturally and work as a backup system to integrate the hemispheres in the human brain. Future studies shall investigate the presence of the TCs in the human brain in different biological contexts and their function and how the TCs are affected by different diseases.

## METHODS

### Animals

All procedures in this study were approved by the Animal Care and Use Committee of the National Institute of Neurological Disorders and Stroke. The marmosets in this study were socially housed with a minimum of two animals per cage. The animals were fed an ad libitum diet and had unrestricted access to water. The marmosets were also provided with toys and other environmental enrichment.

The marmosets used for rsting-state (rsfMRI) were trained for awake fMRI scans(21). Anatomical flexible helmets were designed to perfectly fit the animal’s head and provide robust and comfortable support to minimize head motions during the acquisition. In addition, the animal’s face was monitored with an infrared camera during scans to ensure that the animal was awake and comfortable throughout the session.

### Ex-vivo Diffusion-weighted Magnetic Resonance Imaging

#### Adult Marmosets

The brains of three adult common marmosets (*Callithrix jacchus*) were gadolinium doped(22), placed in a custom-designed 3D-printed holder, and inserted in a 30 mm falcon tube containing Fomblin. Diffusion-weighted images were acquired in a 11.7T/89 mm vertical bore MRI system equipped with a Micro2.5 gradient set of 1500 mT/m and a 30mm quadrature coil. Diffusion-weighted 3-D spin-echo EPI images were acquired using the following parameters: TR/TE = 200/30 ms, δ/Δ = 2.6/5.6 ms, field of view = 35.5 × 25 × 22 mm, matrix = 444 × 312 × 275 yielding an isotropic resolution of 80 μm, number of averages = 1, 240 diffusion-weighting directions split in three shells of 31, 57, and 127 directions, b-values = 1000, 1500, and 2400 s/mm^2^ with 25 b0 images.

### Neonate Marmosets

The brains of two neonate common marmosets (*Callithrix jacchus*) were gadolinium doped (22) and placed in a custom-designed 3D-printed holder containing Fomblin. Diffusion-weighted images were acquired in a 9.4T horizontal bore small animal MRI system equipped with a custom-made 17cm gradient (Resonance Research Inc, Billerica, MA) performing at 450 mT/m gradient strength and a custom-made 30mm coil millipede quadrature coil. Diffusion-weighted 3-D spin-echo EPI images were acquired using the following parameters: TR/TE = 350/42 ms, δ/Δ = 6/14 ms, field of view = 24 × 18 × 16 mm, matrix = 300 × 225 × 200 yielding an isotropic resolution of 80 μm, number of averages = 1, bandwidth = 300 kHz, 90 diffusion-weighting directions split in two shells of 30 and 90 directions, b-values = 1500, 3000 s/mm^2^ with 4 b0 images.

### Cebus

The brain of a Capuchin Monkey (*Cebus apella*) was placed in a custom-designed 3D-printed holder containing Fomblin. Diffusion-weighted images were acquired in a 9.4T horizontal bore small animal MRI system equipped with a custom-made 17 cm gradient (Resonance Research Inc, Billerica, MA) performing at 450 mT/m gradient strength and an 86 mm coil. Diffusion-weighted 3-D spin-echo EPI images were acquired using the following parameters: TR/TE = 250/50 ms, δ/Δ = 6/14 ms, field of view = 48 × 28 × 28 mm, matrix = 240 × 140 × 140 mm yielding an isotropic resolution of 200 μm, number of averages = 1, bandwidth = 300 kHz, 30 diffusion-weighting directions,, b-values = 1500 s/mm^2^ and 60 b-values = 3000 s/mm^2^ with 4 b0 images.

### Macaque

The brain of a Rhesus Macaque (*Macaca mulatta*) was placed in a custom-designed 3D-printed holder containing Fomblin. Diffusion-weighted images were acquired in a 9.4T horizontal bore small animal MRI system equipped with a Bruker BGA20 gradient performing at 300 mT/m gradient strength equipped with an 86 mm coil.

Diffusion-weighted 3-D spin-echo EPI images were acquired using the following parameters: TR/TE = 200/31 ms, δ/Δ = 8/16 ms, field of view = 65 × 55 × 71 mm, matrix = 325 × 275 × 355 yielding an isotropic resolution of 200 μm, number of averages = 1, bandwidth = 300 kHz, 30 diffusion-weighting directions, b-values = 1500 s/mm^2^ and 30 b-values = 3000 s/mm^2^ with 4 b0 images.

### Adult Mice

The brains of six adult C57bl6/J mice were gadolinium doped(18), placed in a custom-designed 3D printed holder, and inserted in a 15mm falcon tube containing Fomblin. Diffusion-weighted images were acquired in a 11.7T/89 mm vertical bore MRI system equipped with a Micro2.5 gradient set of 1500 mT/m and a 15 mm quadrature coil. Diffusion-weighted 3-D spin-echo EPI images were acquired using the following parameters: TR/TE = 350/28 ms, δ/Δ = 2.6/5.6 ms, field of view = 15.36 × 11.52 × 7.68 mm, matrix = 192 × 144 × 96 yielding an isotropic resolution of 80 μm, number of averages = 1, 240 diffusion-weighting directions split in two shells of 30 and 90 directions, b-values = 1500 and 3000 s/mm^2^ with 25 b0 images.

### Neonate Mice

The brains of four neonate C57bl6/J mice were placed in a custom-designed 3D printed holder and inserted in a 10 mm falcon tube containing Fomblin. Diffusion-weighted images were acquired in a 11.7T/89 mm vertical bore MRI system equipped with a Micro2.5 gradient set of 1500 mT/m and a 10mm quadrature coil. Diffusion-weighted 3-D spin-echo EPI images were acquired using the following parameters: TR/TE = 1500/38ms, δ/Δ = 2.6/5.6 ms, field of view = 15 × 9.6 × 9.6 mm, matrix = 300 × 192 × 192 yielding an isotropic resolution of 50 μm, number of averages = 1, 90 diffusion-weighting directions split in two shells of 30 and 60 directions, b-values = 1500 and 3000 s/mm^2^ with 4 b0 images.

### T_2_^*^ anatomical images

The ultra-high-resolution T_2_^*^ image of a common marmoset brain was previously published by Schaeffer et al. (23) and briefly described here. The brain was gadolinium doped (0.2% in PBS) and placed in a custom-designed 3D printed holder containing Fomblin. Flash images were acquired in a 9.4T horizontal bore small animal MRI system equipped with a custom-made 17 cm gradient (Resonance Research Inc, Billerica, MA) performing at 450 mT/m gradient strength and a custom-made 30mm coil millipede quadrature coil. 3-D gradient-echo images were acquired using the following parameters: TR/TE = 100/12 ms, FOV = 35 × 25 × 22 mm, matrix = 1400 × 1000 × 870 yielding an isotropic resolution of 25 μm, number of averages = 1, bandwidth = 50 kHz, flip angle 50.6 (optimized for earns angle at a measured T1=220ms).

### In-vivo Diffusion-weighted Magnetic Resonance Imaging

#### Marmosets

Multishell diffusion MRI data were previously published by Liu 2020 et al. (24) and are briefly described here. Data were collected in a 7T Bruker scanner with a 2D diffusion-weighted spin-echo EPI sequence: TR = 5.1 s, TE = 38 ms, FOV = 36 × 28 mm, matrix size = 72 × 56, slice thickness = 0.5 mm, a total of 400 DWI images for two-phase encodings (blip-up and blip-down) and each has three b values (8 b = 0 s/mm^2^, 64 b =2400 s/mm^2^, and 128 b = 4800 s/mm^2^).

#### Human Ultra-resolution data

The Human in vivo ultra-resolution data was previously published by Wang 2020 et al. (6) and briefly described here. Data were acquired from a single subject across 18 scans with a spatial resolution of 0.76mm 2808 directions split into two different shells (840 b = 1000 s/mm^2^ and 1680 b = 2500 s/mm^2^ and 288 b = 0 s/mm^2^) to ensure accurate shell sampling and SNR at this resolution.

#### Human CCD Subjects

Data from CCD patients were acquired in a Prisma scanner following the Human connectome project protocol (HCP), previously published by Szczupak 2021 et al. (13) and briefly described here. Images were acquired following the HCP criteria for 3.0 Tesla scanners. The diffusion scheme of 200 directions was split into two different shells with b values of 1500 and 3000 s/mm^2^ with a 1.5-mm isotropic resolution acquired in two different phase encoding directions to minimize the drop-out signal.

#### Infant Human Subjects

MR imaging was performed in two infants with non-CCD brain malformations with enlarged ventricles (16 month old (Fig.6G-L): achondroplasia with abnormal gyration in temporal lobes and 10 week old (Fig.6A-F):bilateral frontal and perisylvian polymicrogyria as clinically indicated examinations on a 1.5 T MR system (Siemens Aera) using a head coil. Sedation was used only for the 16-month-old individual, while the 10-week-old infant was imaged without anesthesia. An echo planar diffusion tensor sequence was acquired in an axial plane using 30 gradient-encoding directions, b-values of 0 and 700 s/mm^2^ and a reconstructed asymmetric voxel size of 1.57×1.57×2 mm.

#### Image Denoising

Human images were denoised in Mrtrix (26), while animal images were denoised using the variance stabilization transformation (VST) approach. The noise on the magnitude image is known to follow a Rician distribution (27, 28). Therefore, we applied the forward VST to stabilize the noise variance and convert the signal-dependent noise into signal-independent with a Gaussian-like distribution. Then, the block-matching and 4D filtering (BM4D, a nonlocal transform-domain filter designed for volumetric data corrupted with Gaussian signal-independent noise) algorithm was applied to denoise the images in the VST domain (29). Finally, the inverse VST was used to take data back to its original magnitude space. We emphasize that this approach is particularly advantageous to high-resolution images (30), making it a perfect fit for our high-resolution animal data.

#### Diffusion Imaging Processing

All diffusion images were corrected for eddy-current-induced geometric distortions using FSL software (31), and human images were denoised with Mrtrix software (32) and fitted to the tensor model, visualized, and quantified in DSI Studio software (33). FA values were compared with the non-parametric Kruska-Wallis test.

#### fMRI

Functional MRI data were previously published by Schaeffer at al. (23), and the correlations were obtained through marmosetbrainconnectome.org freely and publicly available resource. To compare the squirrel and rat data to marmoset data, we used fMRI data from our open-access resource (https://marmosetbrainconnectome.org). This database contains resting-state fMRI data from 31 awake marmosets (Callithrix jacchus, 8 females; age: 14–115 months; weight: 240–625 g) that were acquired at the National Institutes of Health (26 animals) and the University of Western Ontario (5 animals).

#### Viral Injection and Histology

For neuronal tract tracing, a 2 μl solution containing 2×10^11^ copies/ μl of AAV9 (AAV9-hSyn-GFP and AAV9-hSyn-mCherry, Penn Vector Core, University of Pennsylvania) was loaded into a 5 μl Hamilton syringe mounted to the arm of a stereotaxic frame (Stoelting Instruments) and injected into the dorsolateral prefrontal cortex (A6DR) of an adult marmoset, and into both ventral and dorsal hippocampus in a separate animal. The stereotaxic coordinates were obtained from the MBM atlas (34). These specific brain regions were chosen because they are known to make corticothalamic interhemispheric connections in different mammalian species (4–7). Four weeks after the injection, the animals were euthanized and perfused with PBS 0.1M followed by PFA 4% in PBS, and their brains were extracted and placed in 30% sucrose in 0.1M PBS for cryoprotection for a week. The brains were coronally sliced into 60μm sections, mounted, and sealed in histological slides. Images were acquired in an AxioVision (Carl Zeiss Microimaging) microscope with a 10x objective lens.

## Funding

This work was supported by the PA Department of Health grant SAP# 4100083102, the National Institute on Aging grants U19 AG074866 and R24 AG073190, Foundation of the State of Rio de Janeiro (FAPERJ), National Council for Scientific and Technological Development (CNPq), as well as by intramural grants from D’Or Institute for Research and Education (IDOR).

## Acknowledgments

We thank the International Research Consortium for the Corpus Callosum and Cerebral Connectivity (IRC5, https://www.irc5.org) members and affiliates for discussions and input. We also thank Professor Kim Phillips, Chair of the Department of Neuroscience at Trinity University, for providing the *Cebus Apella* brain tissue.

## Conflict of interest statement

None declared.

## Notes

### Competing Interest Statement

The authors have declared no competing interest.

