## Supplementary Figure 1 for "Direct interhemispheric cortical communication via thalamic commissures: a new white-matter pathway in the primate brain"

**Supplementary materials**

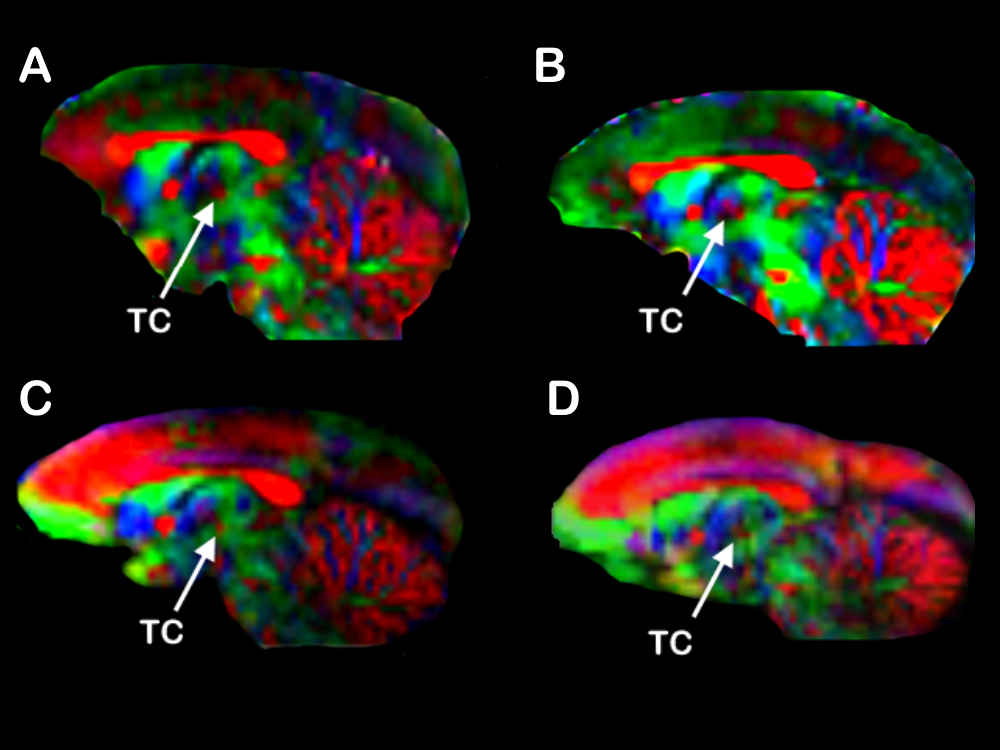

**Supplementary Figure 1. *In vivo* marmoset diffusion image reveals thalamic commissures.** Color-coded diffusion-weighted images show a midsagittal slice of four different anesthetized marmosets. This sequence provides enough spatial resolution (0.5mm isotropic) and fiber discrimination to resolve and clearly identify the Thalamic Commissures (TC). In the Color-coded maps, red represents mediolateral (ML) diffusion, green represents anteroposterior (AP) diffusion, and blue represents dorsoventral (DV) diffusion.

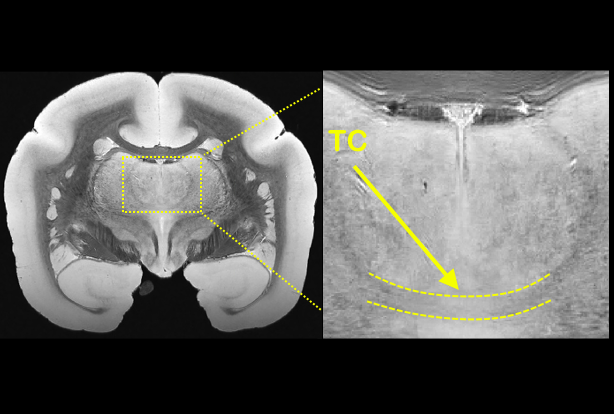

**Supplementary Figure 2. Ultra-high resolution anatomical MRI reveals thalamic commissures.** A coronal view of *ex vivo* imaging of a marmoset brain at 25µm isotropic allows for clear visualization of the commissural pathway through the thalamus (TC) at the level of the nucleus reuniens.

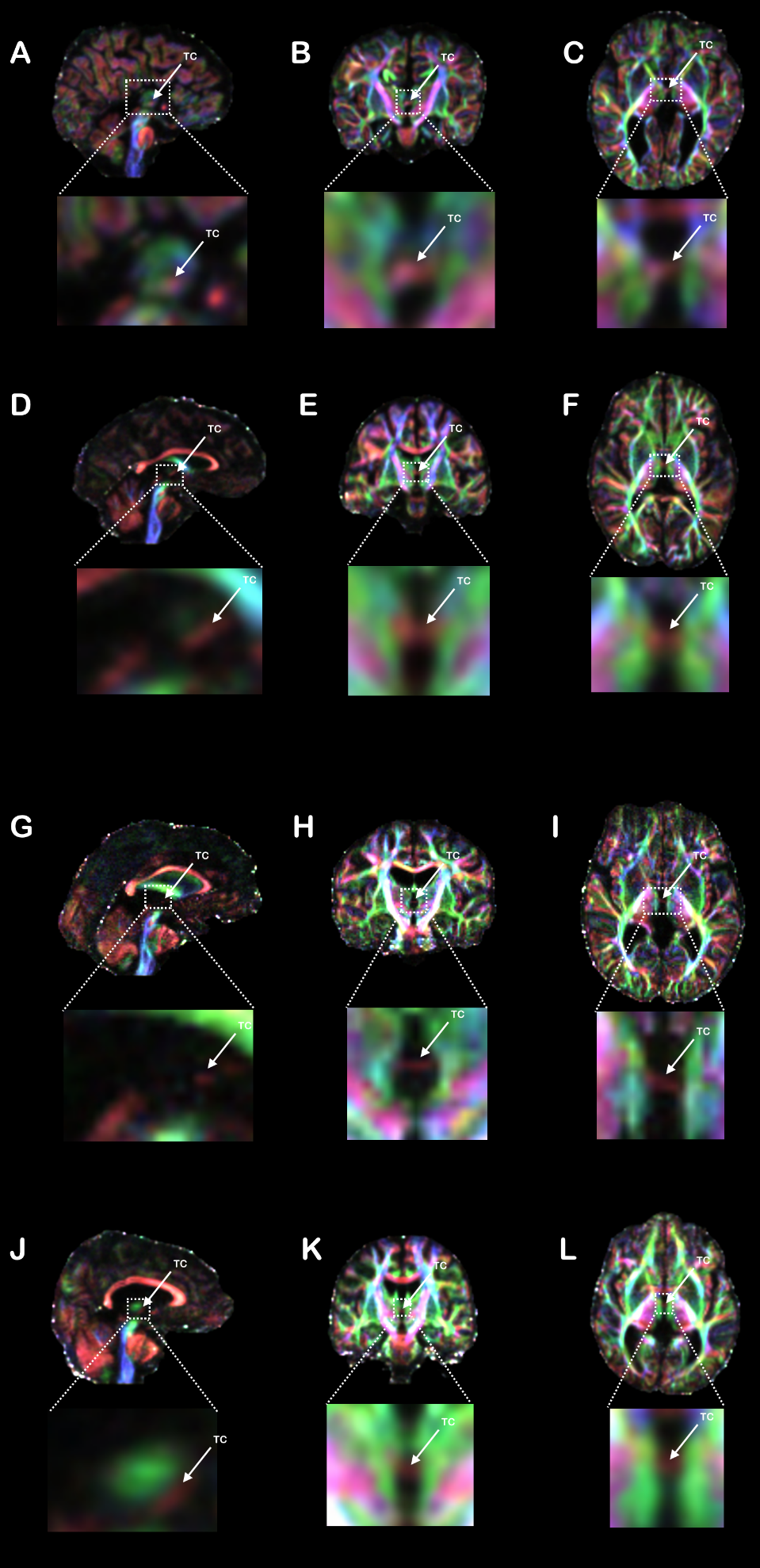

**Supplementary Figure 3. Individual Human callosal dysgenesis subjects presenting callosal dysgenesis.** Color-coded FA maps in the three anatomical planes (sagittal, coronal, and axial) of four human callosal dysgenesis individuals presenting thalamic commissures (TC). In the Color-coded maps, red represents mediolateral (ML) diffusion, green represents anteroposterior (AP) diffusion, and blue represents dorsoventral (DV) diffusion.
